# Local interneurons in the murine visual thalamus have diverse receptive fields and can provide feature selective inhibition to relay cells

**DOI:** 10.1101/2023.08.10.549394

**Authors:** Alexis S. Gorin, Yizhan Miao, Seohee Ahn, Vandana Suresh, Yinan Su, Ulas M. Ciftcioglu, Friedrich T. Sommer, Judith A. Hirsch

## Abstract

By influencing the type and quality of information that relay cells transmit, local interneurons in thalamus have a powerful impact on cortex. To define the sensory features that these inhibitory neurons encode, we mapped receptive fields of optogenetically identified cells in the murine dorsolateral geniculate nucleus. Although few in number, local interneurons had diverse types of receptive fields, like their counterpart relay cells. This result differs markedly from visual cortex, where inhibitory cells are typically less selective than excitatory cells. To explore how thalamic interneurons might converge on relay cells, we took a computational approach. Using an evolutionary algorithm to search through a library of interneuron models generated from our results, we show that aggregated output from different groups of local interneurons can simulate the inhibitory component of the relay cell’s receptive field. Thus, our work provides proof-of-concept that groups of diverse interneurons can supply feature-specific inhibition to relay cells.

## INTRODUCTION

Every relay cell in the dorsal lateral nucleus of the thalamus (dLGN) receives powerful inhibition from local interneurons (Bickford et al., 2010; Montero, 1991). To understand how thalamus processes information, it is imperative to identify the features that these cells encode. In cat and monkey, the few interneurons that have been studied have receptive fields like those of most relay cells—a center surround organization that reflects retinal input (Wang et al., 2011; Wilson, 1989; Wilson et al., 1996). Further, there is a push-pull arrangement of excitation and inhibition within each subregion such that where bright stimuli excite, dark stimuli inhibit and *vice versa* (Wang *et al*., 2011; Wang et al., 2007). It is, thus, easy to imagine that local interneurons supply inhibition to relay cells of the opposite center sign and thus generate “pull”; further, circumstantial evidence supports this arrangement (Martinez et al., 2014; Wang *et al*., 2011; Wilson, 1989; Wilson *et al*., 1996).

Rodents are tasked with navigating local environments very different from those that highly visual animals (e.g., cat, monkey) inhabit. This difference in ecological demand is reflected by disparities in the early visual pathway. For example, retinal ganglion cells in mouse have diverse receptive field types and response properties (Baden et al., 2016; Kerschensteiner and Guido, 2017; Sanes and Masland, 2014). Accordingly, the suite of receptive fields in murine dLGN includes several varieties of On-Off cells (i.e., bright and dark stimuli presented to the same regions of visual space evoke responses of the same sign) (Piscopo et al., 2013) as well center-surround cells (Grubb and Thompson, 2003; Suresh et al., 2016) with push-pull (Suresh *et al*., 2016).

Anatomical differences between mouse and other species are stark. Local interneurons in carnivore (Hamos et al., 1985; Humphrey and Weller, 1988) and primate (Wilson, 1989; Wilson *et al*., 1996) have compact shapes that subtend small regions of dLGN and, hence, retinotopic space. By contrast, dendritic arbors in mouse often traverse most of the nucleus (Charalambakis et al., 2019; Morgan and Lichtman, 2020). Further, the proportion of interneurons in mouse (∼6-10%) (Evangelio et al., 2018; Jager et al., 2021) is less than half of that in carnivore (∼25%) (LeVay and Ferster, 1979) and primate (∼25-35%) (Montero and Zempel, 1986). Last, the number of retinal ganglion cells that converge on single postsynaptic target in dLGN is far greater in mouse (Hammer et al., 2015; Morgan et al., 2016; Rompani et al., 2017) than for highly visual animals (Cleland et al., 1971; Sincich et al., 2007; Usrey et al., 1999). Thus, we were motivated to explore the visual physiology of local interneurons within the murine dLGN.

Recording from local interneurons is challenging. They are small and scarce (Charalambakis *et al*., 2019; Evangelio *et al*., 2018; Jager *et al*., 2021; LeVay and Ferster, 1979; Montero and Zempel, 1986) and cannot be distinguished by standard methods of extracellular recording (Pape and McCormick, 1995; Wang *et al*., 2011). Thus, we took advantage of optogenetic approaches that mouse offers to identify interneurons and relay cells in dLGN *in vivo* and compare the receptive field structures of these two types of neurons. Since most interneurons in mouse primary visual cortex are not selective for stimulus polarity (Liu et al., 2009) or other properties (Kerlin et al., 2010), we expected to find a comparable scenario in dLGN. However, this was not the case. Interneurons in murine dLGN had receptive field structures and sizes almost as diverse as those of relay cells, albeit the distribution of receptive field types was different. To ask how these interneurons might provide the types of feature-specific inhibition that we had previously recorded from murine relay cells (Suresh *et al*., 2016), we developed a simple computational framework. This framework provided proof-of-concept that combined input from several different types of interneurons can recapitulate the suppressive component of the relay cell’s receptive field. Thus, despite substantial variations in morphology and receptive field structure across species, interneurons in dLGN are able to provide precisely tailored inhibition to relay cells.

## METHODS

### EXPERIMENTAL MODELS AND SUBJECT DETAILS

#### Animals

All experiments were performed using mice ≥ 8 weeks old to avoid the visual critical period; there was no detectable difference between males and females so data from both sexes were pooled. Subjects were either Gad2-IRES-cre mice (n=28) ((Taniguchi *et al*., 2011), JAX 010802) or the progeny of these animals crossed with Ai32 mice expressing ChR2(H134R)/EYFP (n=37) ((Madisen *et al*., 2012), JAX 012569). All procedures were approved by the Institutional Animal Care and Use Committees of the University of Southern California following guidelines from the National Institutes of Health.

### METHOD DETAILS

#### Virus expression

Opsins were introduced into GABAergic cells by crossing Gad2-IRES-cre and Ai32 lines or by injection of opsin via adeno-associated viruses (AAV) directly into dLGN of Gad2-IRES-cre mice. To excite interneurons, we introduced channelrhodopsin using: AAV1-EF1a-double floxed-hChR2(H134R)-EYFP-WPRE-HGHpA (Addgene: 20298-AAV1, provided by Karl Deisseroth, ≥ 7 x 10^12^ vg/mL). To inhibit cells, we introduced one of two opsins: AAV1-hSyn1-SIO-stGtACR2-FusionRed (Addgene: 105677-AAV1, provided by Ofer Yizhar (Mahn *et al*., 2018), ≥ 1 x 10^13^ vg/mL) or AAV1-Ef1a-DIO eNpHR 3.0-EYFP (Addgene: 26966-AAV1, provided by Karl Deisseroth, ≥ 1 x 10^13^ vg/mL).

#### Surgical preparation for viral injection

For some experiments, we injected dLGN in Gad2-IRES-cre mice with AAV to restrict expression of opsins to local interneurons. Anesthesia was initiated (2% isoflurane in oxygen 2 L/min) and maintained (1% isoflurane in oxygen 1 L/min). The anesthetized animal was positioned in a stereotaxic device and a small craniotomy made above each dLGN at coordinates −2.2 mm anteroposterior and 2.2 mm lateral relative to bregma. Virus was injected iontophoretically (Nanoject II, Drummond Scientific) via a glass micropipette (tip diameter ∼15-20 μm) lowered 2.5 mm below the brain’s surface (Yu et al., 2016). The pipette was first dipped into a solution containing Fast Green (0.1% w/v) (Sigma-Aldrich) and then introduced into a vial containing 2 μl of AAV solution; after filling the pipette with this mixture, 100nL was ejected over a 5 min period and the pipette withdrawn 10 min later. The scalp was sutured and subjects were treated with ketoprofen and buprenorphine prior to return to the home cage for recovery. Recordings were made three to five weeks post-injection and viral expression was confirmed postmortem by fluorescence microscopy.

#### Surgical preparation for recordings

Prior to electrophysiological recording animals were given an injection of chlorprothixene (5 mg/kg, i.p.) after which anesthesia was initiated and maintained with urethane (0.5-1 g/kg, 10% w/v in saline, i.p.) (Ciftcioglu et al., 2020). The skull was exposed, a small craniotomy made over dLGN, and a headpost affixed. Wound margins were infiltrated with bupivacaine, the brain and eyes kept moist with saline, and body temperature maintained at 37°C.

#### Electrophysiological recording

Whole-cell or cell-attached recordings using pipettes with a potassium-based internal solution were made (Suresh *et al*., 2016) in voltage-clamp mode with a Multiclamp 700B amplifier (Molecular Devices) to damp intrinsic conductances as per previously published protocols (Ciftcioglu *et al*., 2020; Suresh *et al*., 2016) and digitized at 10 kHz with a Power 1401 data acquisition system (Cambridge Electronic Design Ltd). Extracellular recordings were made with a multi-conductor electrode (single-shank, 32-channel, H6b probes (Cambridge NeuroTech) connected to a 32-channel digital multiplexing headstage and a Digital Lynx 4SX-M acquisition system (Neuralynx); high and low pass filters were set to 0.1 Hz and 9 kHz and the sampling rate was 30 kHz.

#### Visual stimulus presentation

Visual stimuli (Figure 1A-I) were generated using a ViSaGe stimulus generator (Cambridge Research Systems) and displayed on a gamma corrected (Dell U2211H LCD) monitor (refresh rate, 70 Hz; stimulus update rate, 28 ms) at a distance of 180 mm, positioned to capture contralateral receptive fields within the appropriate region of visual space. The stimulus display was held at constant luminance (dark, gray, or bright) in order to regulate overall levels of excitability during optogenetic stimulation. The sparse noise stimulus consisted of bright and dark squares, 2°-30° shown at half or full contrast, 16-20 times each in pseudorandom order on a 16×16 grid (grid spacing, 5°) (Ciftcioglu *et al*., 2020; Jones and Palmer, 1987; Suresh *et al*., 2016).

**Figure 1.**
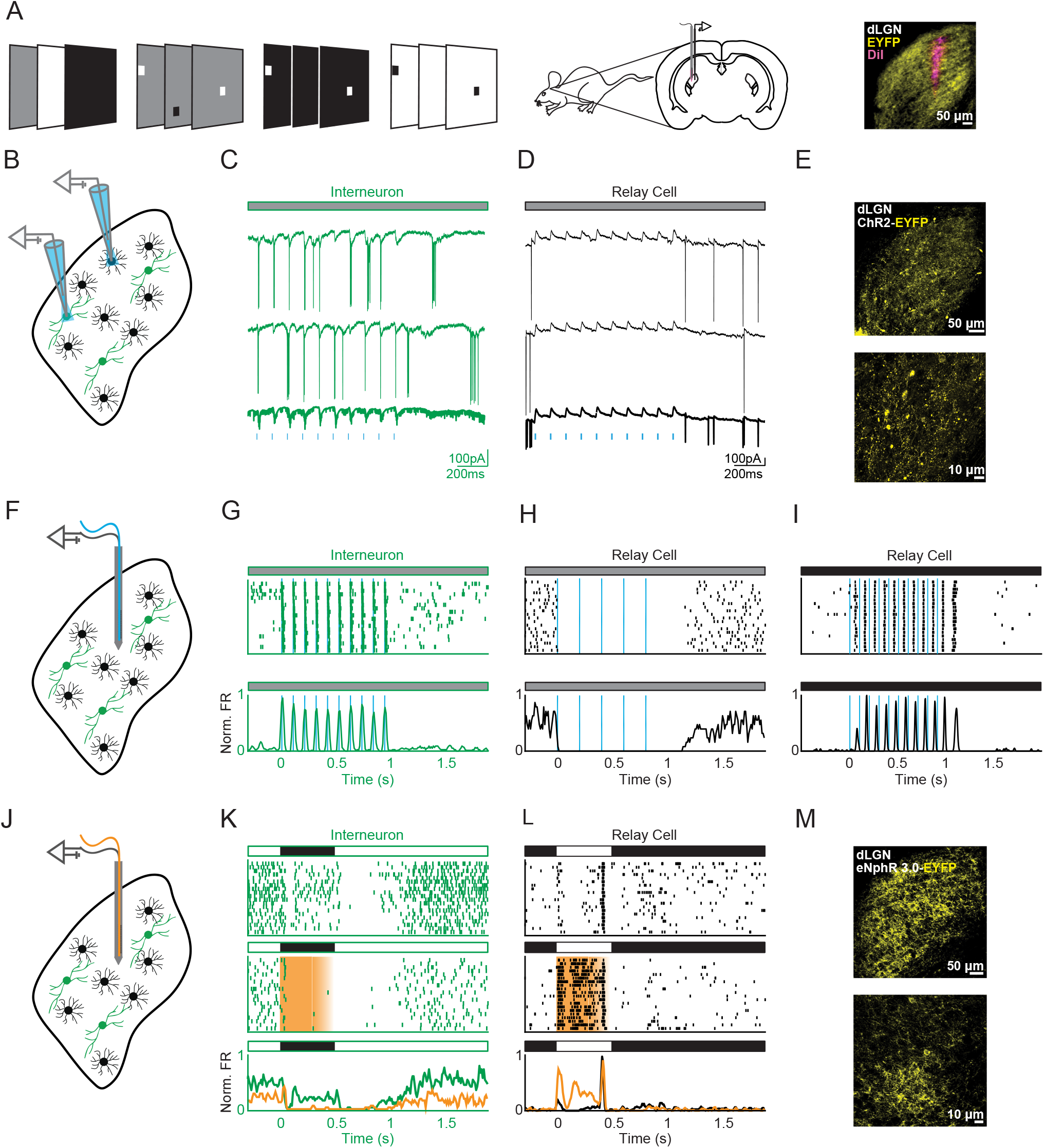
Identifying cell types using optogenetic approaches in mouse dLGN. (A) (from left to right) Different types of stimuli (full field or sparse noise at different contrasts) were shown to mice while recording from dLGN; image of an electrode track (DiI, pink) through tissue in which interneurons express an opsin and reporter (EYFP, yellow). (B) Whole-cell recording from dLGN in which interneuron express ChR2. (C) Pulses of blue light evoked inward currents capped by spikes (two individual traces are shown above the trial average) for an optogenetically labeled interneuron and (D) outward currents for a relay cell. (E) Images of interneurons in dLGN that express ChR2-EYFP. (F) Multielectrode recording from dLGN in which interneurons express the depolarizing opsin, ChR2-EYFP. (G) Pulses of blue light drive spikes at short latency from interneurons, and evoke either (H) complete suppression or (I) silent periods followed by rebound bursts. (J) Multielectrode recording from dLGN in which interneurons express the hyperpolarizing opsin, eNphR3.0-EYFP. (K) Orange light suppressed visually evoked activity for interneurons and (L) disinhibited relay cells. (M) Images of interneurons that express eNphR3.0-EYFP. Banners above panels that display neural recordings indicate when stimulus was white, gray or black.

#### Activation of opsins

For whole-cell and cell-attached recordings, we used an Optopatcher (A-M Systems) (Katz et al., 2013), in which an optical cannula threaded through the pipette delivered light to recording sites, or a standard pipette holder with a separate optical cannula placed nearby. For multi-channel recordings, a flat tip or Lambda-B optical fiber (100-core; 1.2 mm taper-tip) (Cambridge NeuroTech) was fixed to the multi-electrode to guide light to the tip of each shank. The LED light source was either an LSD-1 (A-M Systems) or Optogenetics-LED-Dual (Prizmatix) that was paired with a 200 μm fiber patch cable to a single cannula. Channelrhodopsin and stGtACR2 were activated with 450 nm light and eNphR3 with 650 nm light. Light from the fiber cannula was blocked to prevent spread to either eye. LED pulse length ranged from 5-250 ms and pulse rate varied between 1-10 Hz (Figure 1B-I).

#### Histological confirmation of viral expression and reconstruction of recording sites

For mice injected with AAV, we confirmed that expression of the opsin filled the dLGN. At the end of each experiment, the animal was perfused with 3% paraformaldehyde; the brain was removed, placed in phosphate buffer with 30% sucrose, and cut in 100 μm coronal sections with a microtome. Tissue sections were mounted using ProLong Glass Antifade Mountant (ThermoFisher) and viewed with a standard fluorescent (Zeiss) or confocal microscope (Zeiss). Micrographs were processed with FIJI (Schindelin *et al*., 2012).

We estimated recording sites using complementary methods. For some experiments, multiconductor electrodes were coated with a yellow or a red-shifted lipophilic dye (DiI, ThermoFisher C7000, and DiD, ThermoFisher D12730, respectively) to mark recording sites; uncoated electrodes often left a visible trace in brains prepared for standard light microscopy. Thus, it was possible to view the track created by the electrode (at least its thickest component) in histological sections postmortem (Figure 1A, right). We estimated anteroposterior and mediolateral position by comparing the sections containing tracks with the Allen Brain Reference Atlas. Tracks that crossed more than one section were aligned using Neurolucida (MBF Science). When it was not possible to visualize the track to the tip, we estimated its trajectory by calculating best fit using the ‘allenCCF’ software package (Shamash *et al*., 2018), which also helped register the relative positions of each conductor. Establishing the dorsoventral coordinate for a recording based solely on depth measurement is always subject to error introduced by, for example, variation among brains with respect to atlas coordinates, determining the “zero point” at the brain surface, slight head or electrode tilt, etc. Thus, when possible, we used various additional measures (e.g., relative depth of first and last visual response or other sensory cues) to estimate dorsoventral position. Estimates nearer the top or bottom of the dLGN allowed us to place the recording site in core or shell; for values in between we made no firm assumptions.

### QUANTIFICATION AND STATISTICAL ANALYSIS

#### Optogenetic identification of interneurons and relay cells

For animals in which interneurons expressed channelrhodopsin, cells were classified as interneurons if LED pulses evoked spikes within 10 ms (Figure 1C, G) or as relay cells if firing was suppressed (Figure 1H). Whole-cell recordings confirmed that LED-driven spikes were generated by depolarizing currents and that the prolonged suppression of firing resulted from lasting hyperpolarization (Figure 1H-I). When using an inhibitory opsin, it was important to regulate activity levels so that we could observe suppression as well as excitation. Thus, we used a sequence of full-field luminance epochs (bright, dark, bright or dark, bright, dark) to assess optogenetic modulation of the visually evoked response. The LED was switched on at the start of the central luminance epoch and, after 70% of the duration of this epoch elapsed, ramped down to reach zero power 150 ms after the start of the last luminance epoch. The slow decline (ramping down) of LED power helped avoid the rebound excitation that some opsins cause (Mahn et al., 2016; Raimondo et al., 2012). Cells whose visual responses were suppressed by the LED were classified as interneurons and those whose activity increased were classified as relay cells. (Figures 1J-M).

#### Event detection and sorting

Neural events (i.e., EPSCs and spikes) were detected in whole-cell recordings of membrane currents via techniques we developed previously (Ciftcioglu *et al*., 2020; Suresh *et al*., 2016; Wang *et al*., 2011; Wang *et al*., 2007). Briefly, we applied an adaptive threshold to the first derivative of the intracellular signal such that the smallest potential events included both EPSCs and noise. These events were then sorted using a support vector machine (SVM) (Chang and Lin, 2013) trained with randomly selected events that were manually labeled as EPSCs or noise. Because events near the decision boundary are prone to misclassification, we labeled these manually for additional training and then reclassified the dataset. Finally, spikes were sorted from the EPSCs by repeating the process with SVM-classified neural events.

We used Kilosort2 (Pachitariu *et al*., 2016) to sort spikes recorded with multichannel electrodes into clusters that we manually evaluated with the visualization tool Phy (Rossant *et al*., 2016) to determine if single clusters represented activity from a one neuron alone or, conversely, if different clusters should be merged. Each cluster was associated with the conductor at which spike amplitude was largest. In some cases, it was necessary to remove the LED-induced artifact; we used a local polynomial approximation algorithm (SALPA) for this purpose in order to avoid distorting individual spike waveforms (Wagenaar and Potter, 2002).

#### Receptive field mapping

Spatiotemporal receptive fields were estimated from responses to sparse noise by calculating the spike-triggered average (STA) of the stimulus ensemble (Schwartz et al., 2006); we limited the analysis to 300 milliseconds (t = −300ms) prior to a spike. Receptive fields were qualitatively grouped as center-surround (On center + Off surround or Off center + On surround), On, Off, or On-Off based on responses to the smallest stimulus size that elicited a consistent response.

##### Determining receptive field size

Receptive field size was determined with standard techniques (Wang *et al*., 2007). We fit the initial peak of the STA for each cell with a 2D Gaussian function and then used the average of the semi-major and semi-minor axes of the 1σ contour to calculate response area. For receptive fields with two subregions, we report the size of the largest one.

##### Time course of the On and Off responses

We calculated the temporal STA (tSTA) from the region within the 1σ contour of the 2D Gaussian fitted to the peak of each subregion, after subtracting the mean luminance. The tSTAs were normalized to the absolute maximum (On or Off) for a given cell and the significance criterion for each time point was met if the z-scored first-derivative of the tSTA crossed a threshold ± 0.75 SD from the mean based on the following equation:

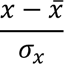

where x is the first-derivative of the tSTA and σ_x_ is the standard deviation of x.

tSTAs divided into two different epochs, or phases. For On and Off cells, the tSTA included an initial “triggering phase” that corresponds to the stimulus that immediately evokes a spike and, often, a “priming phase” of the opposite sign that equates with the stimulus that primes, or facilitates, spiking. Thus, these tSTAs had a peak and a trough. On-Off cells typically had a triggering phase preceded by a priming phase of the same sign to yield a tSTA with two peaks.

The latency of the triggering phase was taken as the first significant time point closest to the spike. The priming phase began at the first significant zero-crossing for On and Off cells, or after the point at which the triggering phase returned to (or near) zero for On-Off cells. The priming phase ended with a return to or near baseline and its strength was defined as its integral.

##### Spike-based Bright-Dark Polarity Score

To assess the relative strength of responses to bright and dark stimuli for a given cell, we adapted an index originally designed for receptive fields constructed from whole-cell recordings of the membrane current (Suresh *et al*., 2016) for use with spikes. This Bright-Dark Polarity Score (Ω) is defined as follows:

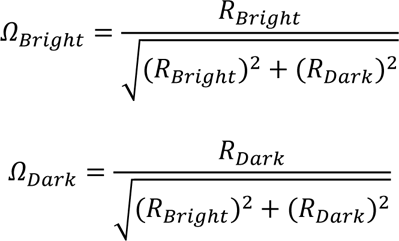

where R_Bright_ and R_Dark_ is the value of the On or Off tSTA at its triggering latency. Ω_Bright_ and Ω_Dark_ values lie on a unit circle. When bright and dark spots evoke responses of the same sign, either excitatory or suppressive, Ω_Bright_ and Ω_Dark_ have the same polarity and so the index values occupy the first or third quadrants of the circle. If bright and dark spots evoke responses of the opposite sign, Ω_Bright_ and Ω_Dark_ have opposite polarities and the index values occupy the second or fourth quadrants of the unit circle. To provide a single number for each score, we converted the bright and dark indices from Cartesian coordinates to polar coordinates and extracted the angle (Θ): −180° < Θ ≤ −90° indicates suppression by both bright and dark; −90° < Θ ≤ 0°, excitation by bright and suppression by dark; 0° < Θ < 90°, excitation by both bright and dark; 90° ≤ Θ ≤ 180°, excitation by dark and suppression by bright.

##### Characterizing spatiotemporal responses of interneurons

We took a model-based approach to ask how spatiotemporal patterns of inhibition we observed in relay cells might arise from the output of the types of interneurons we sampled; these analyses were implemented in Julia (Bezanson *et al*., 2017). First, we used standard linear-nonlinear-Poisson (LNP) models (Chichilnisky, 2001) to describe the spatiotemporal response of On and Off interneurons to sparse-noise stimuli of different sizes and at half contrast:

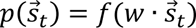

where *p* is probability, 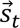 is the vector of stimulus ensemble at a given time, *t*, *f* is the nonlinear filter, and *w* is the linear filter for the space-time separable kernel. For On-Off cells, we modified the model such that two linear kernels and nonlinear filters are used to capture On and Off responses separately:

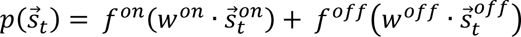

Conventions as above, where 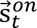 and 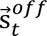 are the ON-only and OFF-only components of the stimulus ensemble vector, 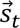.

We fitted the linear component of the model with, at minimum, the PSTH computed from one complete sparse-noise recording, reserving 20% of the dataset to test the model. If multiple recordings with the same sparse-noise sequence were made, all PSTHs were concatenated after each PSTH was normalized to 90% of its maximum. The static nonlinearity was fit with a soft-plus function and the temporal resolution of the model was the stimulus update rate, 28 ms. Model performance was assessed using Pearson’s correlation coefficient, r.

##### Isolating net inhibition from relay cells

To isolate inhibition from whole-cell recordings of the membrane current, we removed spikes and EPSCs from the raw signal. Removing spikes was easily accomplished (MATLAB: medfilt2 function) (Suresh *et al*., 2016; Wang *et al*., 2007). Removing EPSCs was challenging because they vary in size and shape and often overlap. We detected EPSCs as described in **Event detection and sorting** and divided them into clusters based on amplitude (area under the peak of the first derivative of the EPSC) and slope (the peak value of the first derivative of the EPSCs), as we had done earlier (Suresh *et al*., 2016). These clusters contained EPSCs of similar size and shape and, thus, allowed us to fit templates (a linear rise, exponential decay function (Wang *et al*., 2007)) to the cluster mean. We then used each EPSC time (*t*_*ij*_) and corresponding template 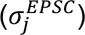 to form a signal that represented contributions of EPSCs to the raw membrane current:

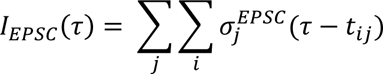

where *t*_*ij*_ is the time of the *i*th EPSC of the *j*th cluster.

Subtracting the spike and template EPSC trains from the raw membrane current yielded a waveform dominated by slow outward currents— net inhibition. We downsampled this residual trace to the stimulus update rate, rectified it to remove uncaptured excitatory currents, and then used standard methods of reverse correlation (Schwartz *et al*., 2006; Wang *et al*., 2007) to compute the suppressive spatiotemporal receptive field. To increase the signal to noise ratio of the suppressive spatiotemporal receptive fields, we fit them with LN models, using a manually chosen L1 regularization term for each residual trace and a rectified linear nonlinearity (Suresh *et al*., 2016; Vaingankar et al., 2012; Wang *et al*., 2011).

##### Building interneuron ensembles to form the suppressive receptive field

To increase the pool of available interneurons for our simulation of the suppressive field, we generated a large library by translating the modeled receptive fields in space and shifting them slightly in time (e.g., an earlier or later stimulus frame). We then used the genetic algorithm to select the (modeled) interneuron that provided the largest contribution to the suppressive field and iterated the process (maximum 20 interneurons). Inputs with negative weights were excluded as these, in essence, provide disinhibition. The output of the interneuron ensemble was compared to the corresponding suppressive field using Pearson’s correlation coefficient.

#### Statistics

All statistics were calculated using MATLAB (Mathworks). When two populations were compared, the distribution of data from each group was tested for normality (Shapiro-Wilk test; null hypothesis: normal distribution), and for equal variance (two-sample F-test; null hypothesis: both distributions are normal and have equivalent variance (MATLAB function ‘vartest2’)). If both conditions were met, we used a two-sample t-test (MATLAB function ‘ttest2’). If only the first condition was met, we used the two-sample Welch’s t-test (MATLAB function ‘ttest2’). For non-parametric datasets, we used the Wilcoxon rank sum test (MATLAB function ‘ranksum’) in addition to the two-sample Kolmogorov-Smirnov test (MATLAB function ‘kstest2’). All statistical tests were two-tailed with sample size indicated in each figure and/or legend. Significance levels were indicated by asterisks: n.s.: p>=0.05; *: p<0.05; **: p<0.01; ***: p<0.001. Last, statistical variation for a given data set is reported as mean ± SD.

## RESULTS

Our goal is to understand how local interneurons in dLGN contribute to receptive field structures and response properties of relay cells. Towards this end, we compared visually evoked responses from optogenetically identified interneurons (n=27) and relay cells (n=127) in mouse dLGN (Figure 1A) (expression was driven by viral vectors (n=32) or genetically (n=122)) and used computational approaches to determine how input from interneurons in our dataset might give rise to patterns of inhibition recorded from relay cells (Suresh *et al*., 2016). Our sample was obtained from 65 adult mice of both sexes. Recordings from interneurons were distributed throughout dLGN (−1.8 to 2.5mm anteroposterior and 2.0 to 3.3mm mediolateral) based on alignment with the Allen Brain Reference Atlas; 10 interneurons were clearly in core, 2 in shell and the remaining 15 nearer presumed borders between regions; relay cells were sampled in the same territory, but more densely.

### Recording visual responses from optogenetically identified interneurons and relay cells

Because sampling interneurons *in vivo* is challenging, we used an optogenetic approach to distinguish between different types of cells (Lima et al., 2009) (Figure 1A). We used the Gad2-cre mouse line to drive expression of various opsins including Channelrhodopsin-2 (ChR2), halorhodopsin (eNphR3.0), and soma-targeted anion-conducting channelrhodopsin (stGtACR2), either by crossing this line with the Ai32 line or injecting viral vectors in dLGN.

For assurance that ChR2-mediated changes in spike rate reflected changes in membrane currents, we began our studies with whole-cell recordings (Figure 1B). When pulses of blue light delivered through a cannula in the dLGN evoked depolarizations (often capped by spikes), we classified the cell as an interneuron (Figure 1C). If, instead, the LED evoked hyperpolarizing responses (presumably generated by activating presynaptic interneurons), we classified the neuron as a relay cell (Figure 1D). Each panel shows two individual sequences of a pulse train above the average of all trials (bold); recordings were made while the animal viewed a full-field gray stimulus (banner at top). Expression of ChR2 was evident in interneurons (Figure 1E). The same optogenetic protocols during extracellular recordings (Figure 1F) evoked volleys of spikes at short latency for interneurons (Figure 1G) and either prolonged suppression (Figure 1H) or silent periods followed by bursts of spikes (presumably T-type calcium channel mediated bursts (Llinás and Jahnsen, 1982)) for relay cells, as illustrated by raster plots placed above peristimulus time histograms (PSTHs) (Figure 1I).

We complemented experiments with ChR2 by using inhibitory opsins, such as eNphR3.0 (Figure 1J). To assess optogenetic modulation of activity during these experiments, it was often necessary to increase neural firing rate with a visual stimulus. Cells whose visually evoked activity was reduced by photostimulation were classified as interneurons (Figure 1K) and those for which suppression was relieved were classified as relay cells (Figure 1L). We confirmed virus expression histologically (Figure 1M). These optogenetic techniques thus provided a means to classify cells in our preparation.

### Receptive field structures of interneurons are nearly as diverse as those of relay cells

Unlike cat, for which almost all relay cells have center-surround receptive fields and push-pull responses (Hubel and Wiesel, 1961; Wang *et al*., 2007), there is great diversity in mouse (Durand et al., 2016; Marshel et al., 2012; Piscopo *et al*., 2013; Scholl et al., 2013; Suresh *et al*., 2016; Zhao et al., 2013). Although many relay cells in murine dLGN have receptive fields that resemble those in cat, the rest of the population includes various types of On-Off (On and Off subregions overlap) receptive fields (Suresh *et al*., 2016). Thus, we asked if this variety of receptive field structures were mirrored by local interneurons, or if they, like most GABAergic cells in murine visual cortex (Kerlin *et al*., 2010; Liu *et al*., 2009) supply nonspecific inhibition to their excitatory counterparts.

Our finding was that interneurons had diverse receptive field structures (Figure 2). We mapped receptive fields with sparse noise, sequences of individually flashed bright and dark squares (Suresh *et al*., 2016), so that we could separate the contributions of the On and Off responses (see **Methods**) and depicted receptive fields as contour plots made from the peak frame of the spike-triggered average of the stimulus ensemble (STA). The most common types of receptive fields we recorded were On-Off (Figure 2A, left); maps constructed from responses to bright (On) stimuli border those made from responses to dark (Off) ones. These fields ranged from large and patchy to small and discrete. STAs for On (Figure 2A, middle) and Off (Figure 2A, right) cells spanned a similar range in size and shape. Finally, a subset of On and Off cells had a surround and, thus, assumed a center-surround profile (Figure 2B). Plotting the distribution of receptive field types (qualitatively determined based on the smallest sparse-noise stimulus that elicited a robust response) for interneurons alongside that for relay cells revealed differences between the two populations (Figure 2C). Notably, the proportion of interneurons (26%) with center-surround receptive fields was small compared to that for relay cells (56%). Commensurately, there were roughly double the proportion of interneurons with On-Off receptive fields. We have yet to find subtypes defined by exceptionally strong inhibition (i.e., Suppressed-by-Contrast (SBC) cells, for which both bright and dark stimuli evoke strong inhibition, and W3 cells, for which all but small stimuli evoke strong suppression (Piscopo *et al*., 2013; Suresh *et al*., 2016)); these are rare in dLGN, however.

**Figure 2.**
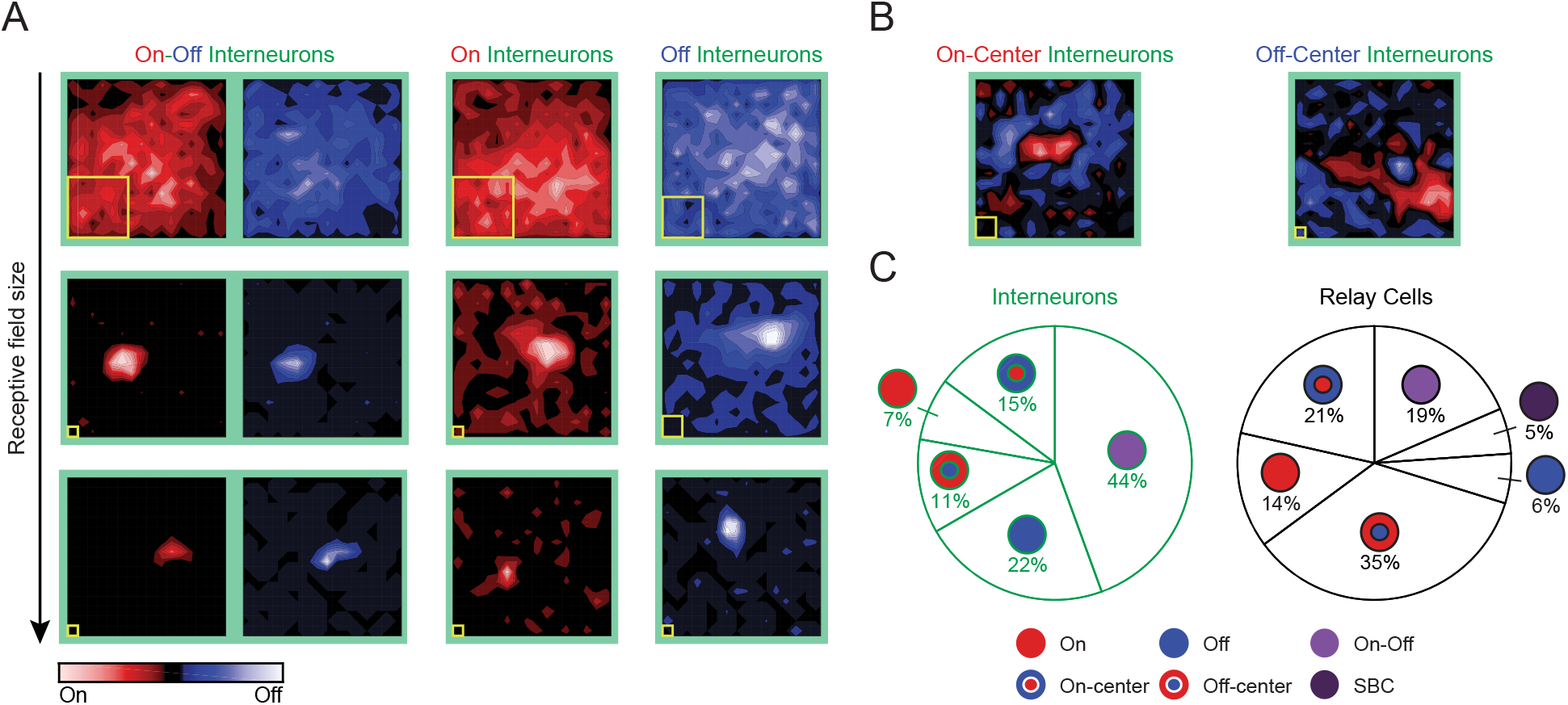
Receptive fields of interneurons are diverse. (A) Receptive fields of On-Off, On, and Off interneurons are shown as contour plots made from responses to dark (blue) or bright (red) sparse-noise stimuli; yellow squares indicate stimulus size and the spacing of the stimulus grid was 5°. Receptive fields within class are organized from large to small as indicated by the arrow at left; note large stimuli were often necessary to evoke responses from cells with large receptive fields. For On-Off cells, bright and dark contour plots are arranged side by side while only bright maps are shown for On cells and only dark maps for Off cells. (B) Receptive fields of On-center and Off-center interneurons; here the contour plots were made by subtracting the dark from the bright maps. (C) Pie-charts show qualitative distributions of the different types of receptive fields of interneurons (n=27) and relay cells (n=205).

### The distributions of receptive field sizes for interneurons and relay cells are similar

Past results from carnivore (Martinez *et al*., 2014) and primate (Wilson, 1989; Wilson *et al*., 1996) show that receptive fields of interneurons are larger than those of neighboring relay cells, a result consistent with the relative frequency of the two cell types (LeVay and Ferster; Montero and Zempel, 1986). Given that the proportion of interneurons is even smaller in mouse than carnivore and primate (Evangelio *et al*., 2018; LeVay and Ferster; Montero and Zempel, 1986) and the extent of dendritic arbors with respect to the retinotopic map far greater (Morgan and Lichtman, 2020; Seabrook et al., 2013), it might seem as if the receptive fields of murine interneurons should be markedly larger than those of relay cells (with the caveat that some relay cells in mouse dLGN have large receptive fields themselves) (Ciftcioglu *et al*., 2020; Kerschensteiner and Guido, 2017; Suresh *et al*., 2016). To compare the sizes of the receptive fields of relay cells and interneurons, we used the 1σ contour of the 2D Gaussian fit to the peak frame of the STA. The distribution of receptive field sizes was comparable for both populations (Figure 3A). This overall similarity in response area was consistent across receptive field types (Figure 3B); an apparent trend towards slightly larger fields for interneurons did not reach statistical significance.

**Figure 3.**
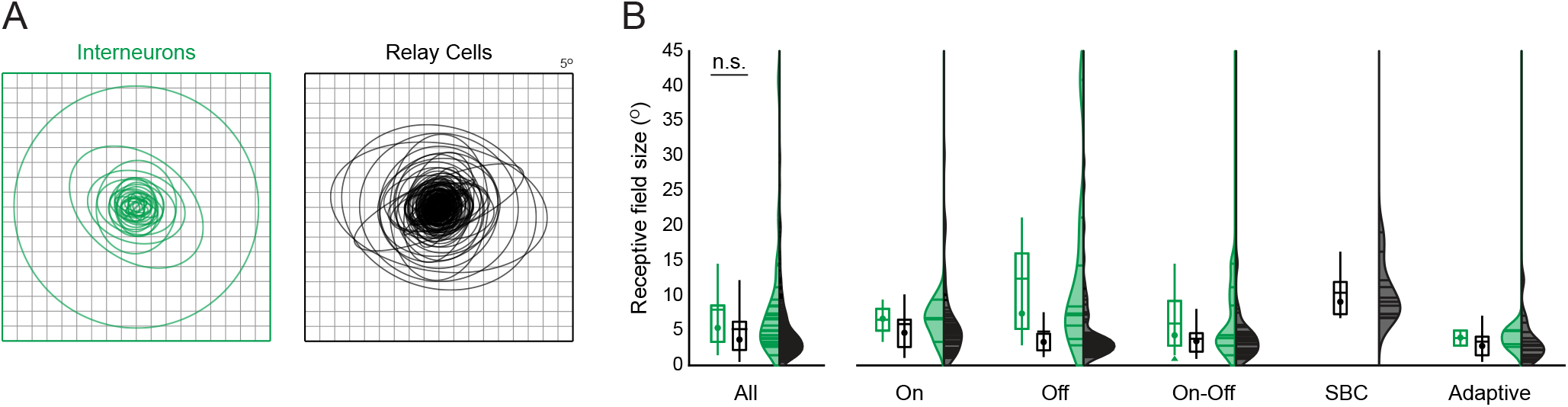
The range in receptive-field sizes for local interneurons and relay cells is similar. (A) 1σ contours of 2D Gaussian fits to the receptive fields of interneurons (green ellipses) and relay cells (black ellipses) aligned to the center of the stimulus grid (interneurons, n = 27; relay cells, n = 205). Fits were made to the full extent of each (sub)region for On, Off, On-Off cells or to the centers for center-surround cells. (B) Leftmost pair of box and violin plots show that the distributions of receptive field sizes for interneurons (green) and relay cells (black) are statistically indistinguishable; Wilcoxon rank-sum test and two-sample Kolmogorov-Smirnov test (p>0.05). For box plots, horizontal lines indicate the mean, circles denote the median, and vertical lines the SD; for violin plots, horizontal lines are individual values. Remaining plots show distributions of receptive field sizes for individual classes of cell: On, Off, On-Off, SBC, and Adaptive (see Figure 4 for an illustration of adaptive cells).

### Stability of stimulus preference across stimulus strength

So far, the receptive fields we have illustrated were mapped using the smallest effective stimulus size at half (50%) contrast (i.e., black and white squares on a mean gray background); this standardized approach reduced the influence of stimulus strength on the comparison of the visual responses across cell types (Wienbar and Schwartz, 2018). It has become clear, however, that neurons in dLGN receive input from many ganglion cells (Hammer *et al*., 2015; Litvina and Chen, 2017; Morgan *et al*., 2016; Rompani *et al*., 2017), and this led us to ask if stimuli of different strengths would recruit responses with different properties. To address this question, it was useful to quantify receptive field type. Thus, we adapted an index called the Bright-Dark Polarity Score (Suresh *et al*., 2016) to assess the extent to which a cell preferred, or was inhibited by, dark or bright stimuli of a given size or contrast (the index did not take surrounds into account). First, we calculated the temporal STA (tSTA) from the region that fell within the 1σ contour of the 2D Gaussian fit of the initial peak of the receptive field. An idealized tSTA is plotted as normalized stimulus contrast against time (moving backwards from 0 ms, the instant at which each spike occurred) (see **Methods**) (Figure 4A). This curve splits into a triggering phase (the stimulus that immediately evokes a spike; e.g., the STAs in Figure 2 correspond to the peak of this phase) and an earlier priming phase (the stimulus that facilitates firing). The latency and duration of each phase was calculated using time points that deviated significantly (± 0.75 SD) from the mean, as illustrated in Figure 4A. We used the triggering phase to calculate the Bright-Dark Polarity Score and plotted the results along a circle divided into four (color-coded) quadrants that indicate receptive field type (Figure 4B). Positive Bright Scores and/or negative Dark Scores fell in the fourth (lower right) quadrant and were classified as “On”, negative Bright and Dark Scores fell in the third quadrant were classified as SBC, and so on. To describe the position of each response within the circle simply, we calculated the angle (Ɵ) in polar coordinates (Figure 4C; Figure S1A-B).

**Figure 4.**
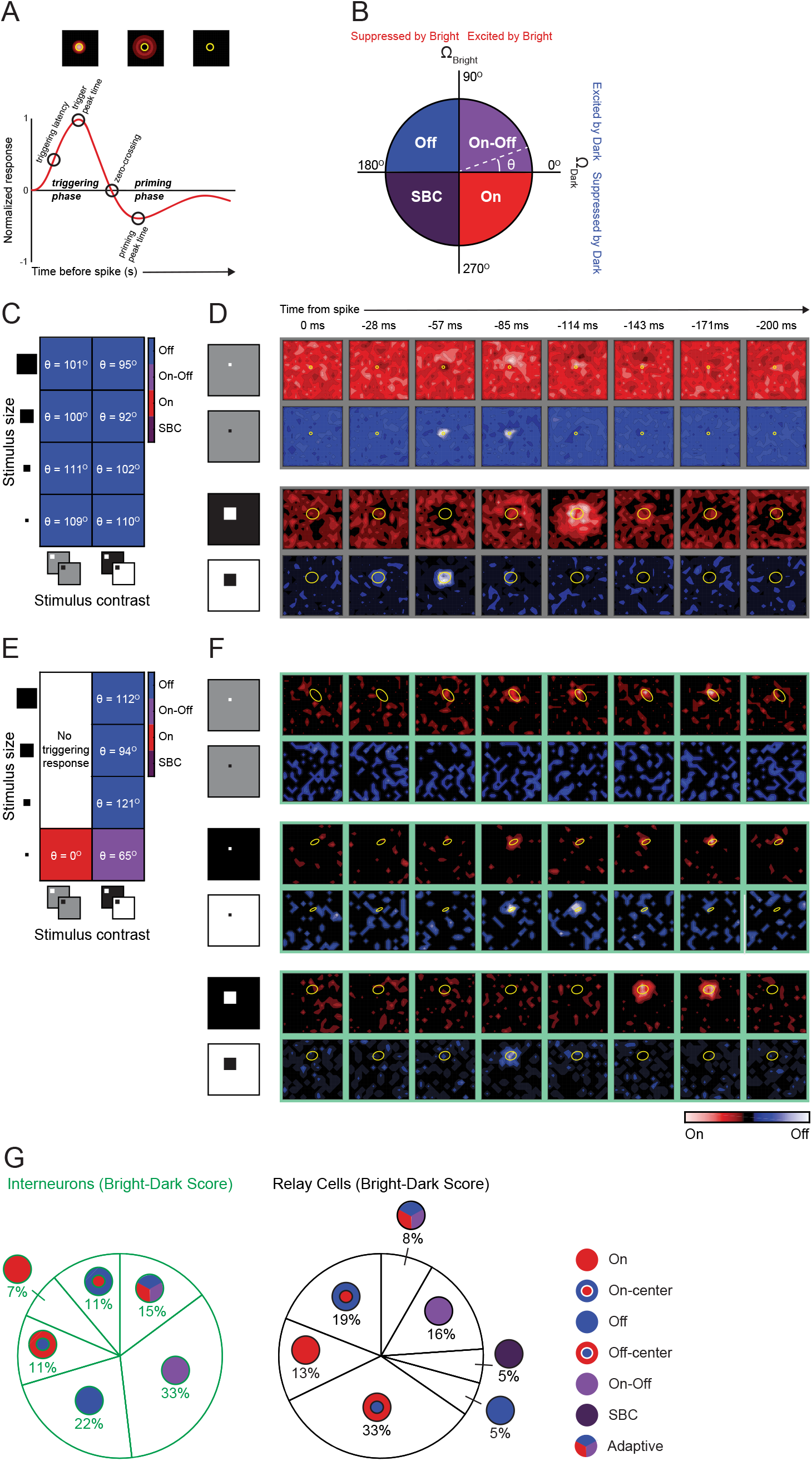
Quantification of receptive field class and constancy across stimulus strength. (A) Graphical explanation of the temporal evolution of the STA (tSTA) shows how a neural response divides into a “triggering phase” that evokes a spike and a preceding “priming” phase and illustrate the peak and time-course of each phase. (B) Values from the Bright-Dark Polarity Score calculated for a given tSTA can be plotted along a circle with four quadrants that define different categories of receptive fields (Off, blue, On-Off, purple, On, red, SBC, deep purple) and can be represented by a single value, θ. (C and D) Results for a cell whose receptive field class (Off) remained constant across stimulus sizes and contrasts, as illustrated with a color-coded grid of Bright-Dark-Polarity Scores calculated from responses to eight stimulus conditions indicated by icons at left and bottom. (D) tSTAs shown as a sequence of frames (yellow overlays are 2D Gaussian fits to the peak frame) for small, low-contrast bright and dark squares shown above those made using large stimuli at high contrast. (E and F) Example of an “adaptive” interneuron whose receptive field class changed as a function of stimulus parameters, conventions as in (C and D). The receptive field was classified as On for small half-contrast stimuli, On-Off for small high-contrast stimuli, and Off for large high-contrast stimuli. (G) Pie-charts of different types of receptive fields of interneurons (n = 27) and relay cells (n = 205) categorized quantitatively by the Bright-Dark Polarity Score (and qualitatively as center-surround) highlight the presence of “adaptive” cells.

For most cells, receptive field type remained the same across stimulus size or contrast, as illustrated for an Off-center On-surround relay cell with push-pull responses (Figures 4C and 4D). tSTAs made for small stimuli displayed at half contrast are shown above those made for full contrast stimuli (Figure 4D, top). Larger and higher contrast stimuli evoked stronger responses but did not change the overall type of the receptive field (Figure 4D, bottom). The Bright-Dark Polarity Score remained similar across stimulus contrasts and sizes (Figure 4C; Figure S1A).

For a subset of cells, receptive field class changed as a function of stimulus parameters (see **Methods** regarding verifying spike clustering). We use the term “adaptive” to describe these cells, with an example neuron illustrated in Figures 4E, 4F, and S1B. Small spots at half contrast evoked a slowly evolving On response but no Off response (Figure 4F, top), while small stimuli at full contrast evoked both On and Off responses (Figure 4F, middle). Large stimuli at full contrast elicited an Off response only (Figure 4F, bottom), and larger spots at half contrast failed to drive the cell (not shown). These shifts in response type are most easily appreciated by viewing the graphical depiction of the Bright-Dark Polarity Score (Figure 4E).

Response dynamics during the priming phase varied so widely from cell to cell that we could not include this interval in our quantitative analysis. Still, we make note of a small population of relay cells and interneurons that were noticeably suppressed by both bright and dark during the priming phase (Figure S1C-I). We do not recall observing such patterns in our previous recordings from cat (Wang et al., 2010; Wang *et al*., 2011; Wang *et al*., 2007).

All told, after analyzing responses for our entire dataset across stimulus contrasts and sizes, we found that 15% of interneurons and 8% of relay cells had adaptive receptive fields. We generated a second set of pie charts to reflect this finding; compare Figure 4G with Figure 2C.

### Reconstructing inhibition in the relay cell’s receptive field

Can the receptive field structures of interneurons that we mapped explain patterns of visually evoked inhibitory currents that we had recorded previously (Suresh *et al*., 2016)? An ideal approach to this question would involve optimizing stimulus parameters for each interneuron and obtaining its response to the identical stimulus set that had been used for whole-cell analyses of relay cells. This approach is not realistic, however, because recording time is often insufficient to display the full range of stimulus sizes at different contrasts. Thus, we developed a computational framework to predict how interneurons would respond to stimuli of a given size or contrast.

We began by building linear-nonlinear Poisson models (LNP) (Chichilnisky, 2001; Paninski, 2004) fitted to visual responses recorded from different types of interneurons. The receptive field of an On or Off cell was described as the product of two linear kernels, one spatial and one temporal (Chichilnisky, 2001), and the nonlinearity was estimated using standard techniques (Karklin and Simoncelli, 2011; Zoltowski and Pillow, 2018). The only nonstandard approach we took was to separate the analysis of On and Off responses for On-Off cells. Figure 5 depicts the spatial and temporal filters for an example On (Figures 5A and 5B), Off (Figures 5B and 5C), and On-Off (Figures 5E and 5F) cell next to biologically-derived (pale green frames) and modeled (dark green frames) STAs. The overall shapes of the STAs computed from recordings of the interneurons were well approximated by the output of the filters estimated by the corresponding models. The modeled STAs also had the benefit of reducing noise. We then expanded the available pool of modeled interneurons by cloning each one into many such that each type of receptive field was represented across visual space and jittered to reflect different response latencies.

**Figure 5.**
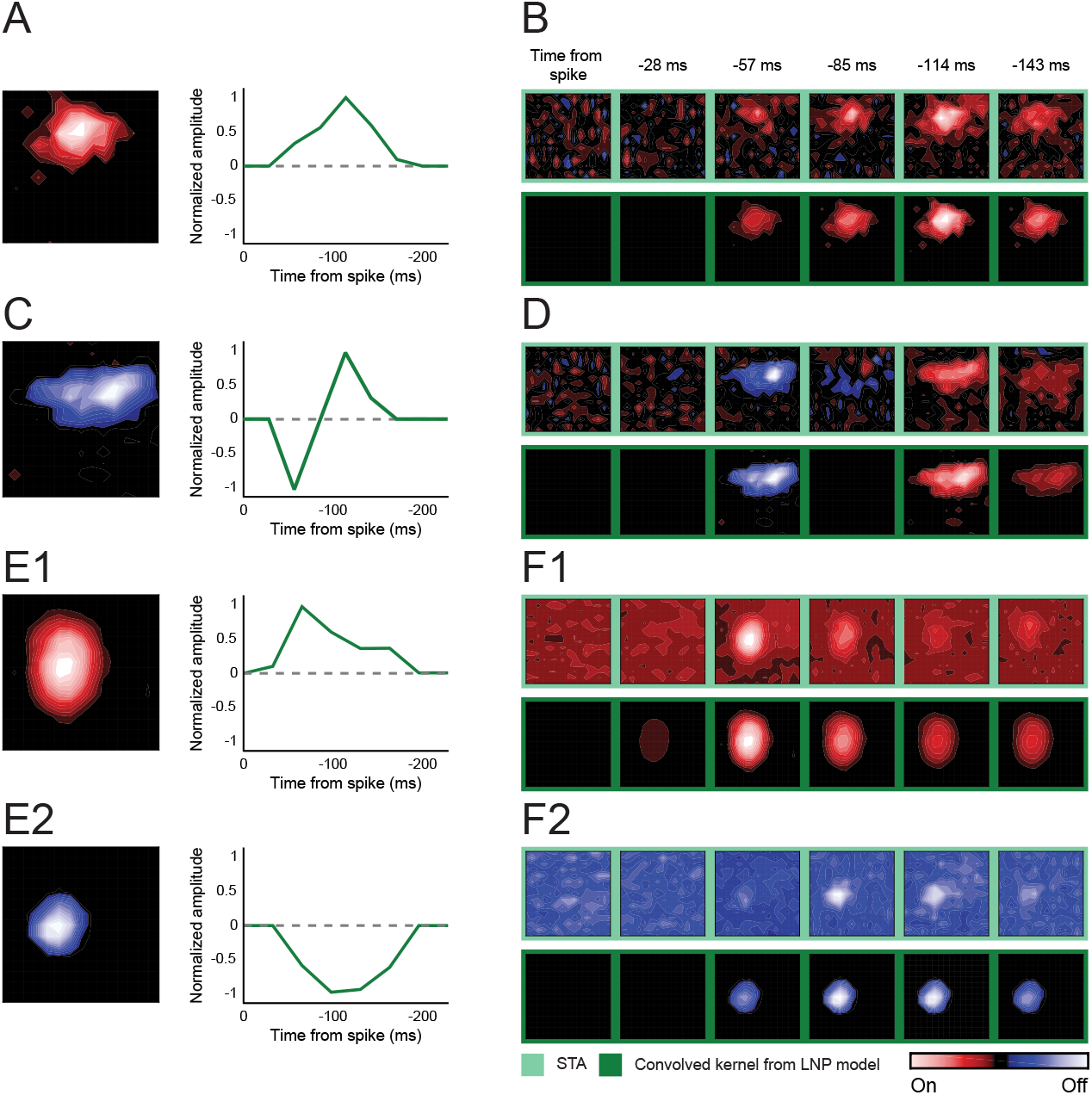
Simulating responses of interneurons with LNP models. (A) Spatial and temporal filters generated from responses of an On interneuron to sparse noise. (B) tSTA estimated from the neural response (top, light green) and the associated LNP model (bottom, dark green). (C and D) Same as A and B but for an Off interneuron. (E1, E2 and F1, F2) For this example On-Off interneuron, spatial and temporal filters and tSTAs computed for bright and dark responses are shown separately; otherwise, conventions as above.

To link the output of the modeled interneurons to postsynaptic inhibition in relay cell, we created a simulation framework. We chose the most common case first, a center-surround relay cell (Figure 6). The receptive field of an On-Center relay cell is shown as a contour plot next to an array of trace pairs in which averaged responses to bright (gray traces) and dark (black traces) squares are organized point by point along the stimulus grid; the center and surround are approximated by dashed contours (Figure 6A). Membrane currents reveal strong push-pull in the center and weaker push-pull in the surround; trace pairs in boxes are shown at increased gain in the insets placed below the contour maps.

**Figure 6.**
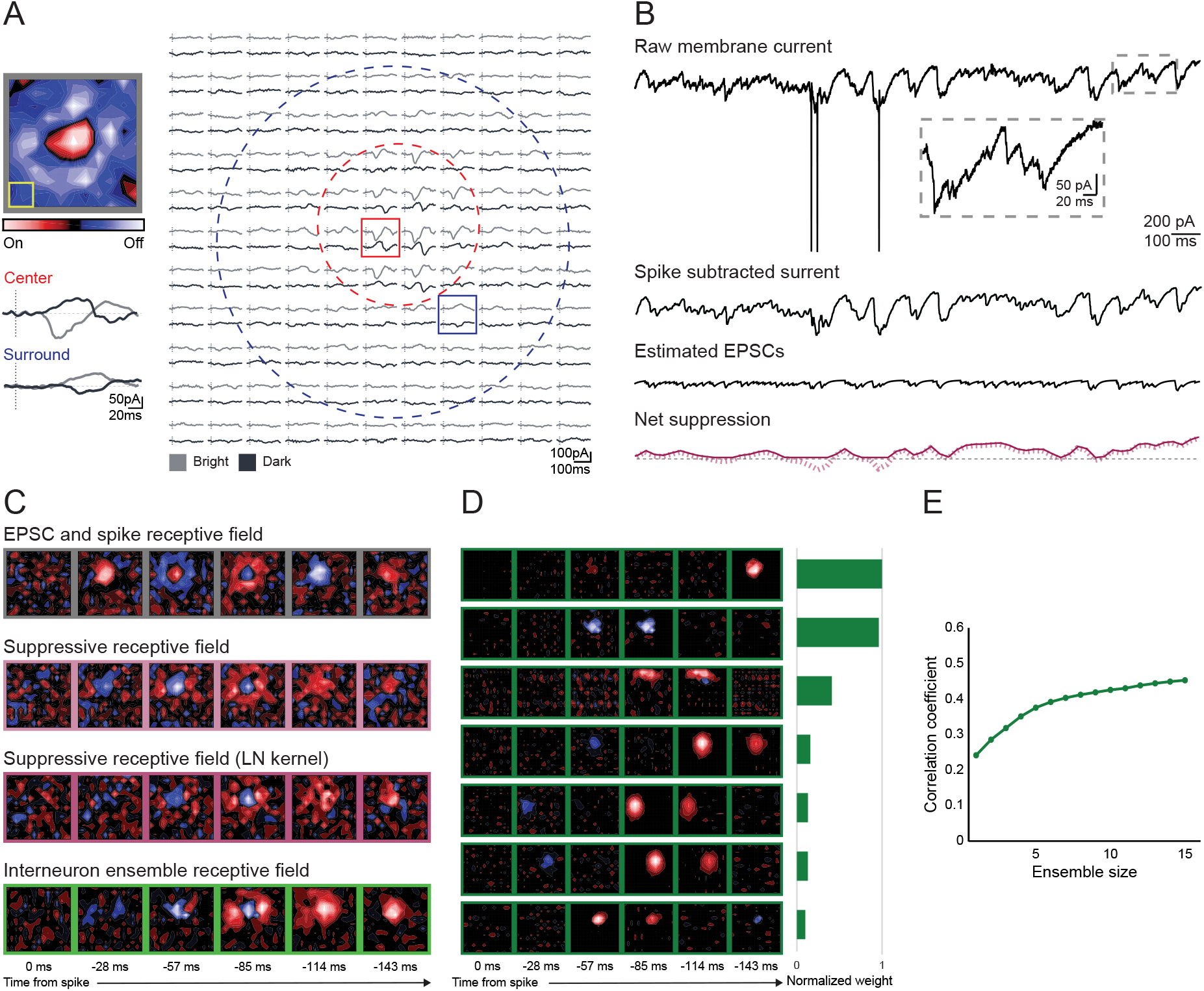
Simulation demonstrating how pooled input from different types of interneurons can generate feature-specific inhibition for an On-center relay cell. (A) Receptive field of an On-center relay cell mapped using sparse noise shown as a contour plot (top left) and as an array of trace pairs (right) in which the averaged responses to bright (gray traces) and dark (black traces) squares are placed at corresponding positions in the stimulus grid, and where dashed outlines approximate the On (red) and Off (blue) subregions; stimulus size 5°. Traces in insets in the center and surround are shown at higher gain to illustrate push-pull responses (bottom left). (B) Continuous recording of the raw membrane current (inset reveals EPSCs at high gain) is separated into three components below. The spike-subtracted current is shown above the EPSC train. These two components are subtracted from the raw current to yield a residual trace (bottom, dashed pink line), which was then rectified to remove uncaptured excitation (solid pink line). (C) Receptive fields made by reverse correlation of excitatory events (EPSCs and spikes) (top) and suppressive currents (the rectified residual) (2nd row) are shown above a linear-nonlinear model of the suppressive field (3rd row) and a simulation (4th row) of that field made by summing input from seven interneurons whose modeled spatiotemporal receptive fields and relative contributions are shown in (D). (E) The correlation coefficient between increasing number of interneurons and the (modeled) suppressive field.

To move forward with our analysis, it was necessary to isolate the underlying net suppressive (outward) currents from the raw membrane currents recorded while the stimulus was displayed (Figure 6B). While we had solved a similar problem for recordings from cat (Wang *et al*., 2007), mouse presented a greater challenge. Retinogeniculate convergence ratios are small in cat (Cleland *et al*., 1971; Usrey *et al*., 1999), but high in mouse (Hammer *et al*., 2015; Maher et al., 2023; Morgan *et al*., 2016; Rompani *et al*., 2017). As a result, EPSCs recorded from murine relay cells vary considerably in size and shape and are often superimposed. Thus, we used a support vector machine (Chang and Lin, 2013; Suresh *et al*., 2016) to classify as many EPSCs as possible (usually the large majority) for a given relay cell and then performed K-means clustering (Hartigan and Wong, 1979) to generate EPSC templates of different sizes. To extract net suppressive currents, we subtracted spikes and EPSCs (represented as their templates) from the raw recording, downsampled this residual signal to the stimulus update rate, and rectified it to remove uncaptured excitation (Figure 6B). The receptive field generated from spikes and EPSCs (Figure 6C, top) had the opposite sign of that generated from the suppressive signal—the suppressive field (Figure 6C, second row); see **Methods.** Thus, we reproduced the basic push-pull pattern of excitation and inhibition within the relay cell’s receptive field.

We used an LN model (Suresh *et al*., 2016; Wang *et al*., 2011) to sharpen the basic shape of the suppressive field (Figure 6C, third row) and proceeded to determine its potential origin. This process was accomplished by using the genetic algorithm to select candidate inputs from our library of interneuron models. For this relay cell, aggregated input from seven interneurons recapitulated the overall structure of suppressive field (Figure 6C, bottom row). Inputs with negative weights were excluded from the fitting process as these, in essence, are disinhibitory. The tSTAs for each interneuron that was chosen to simulate the suppressive field and the relative weights of their contribution are provided in Figure 6D. The correlation between the simulated and biological suppressive fields for increasing numbers of inputs is shown in Figure 6E.

A parallel analysis for an SBC relay cell is provided in Figure 7; conventions are the same as for Figure 6 except that the On and the Off maps for both the relay cell and for the interneurons that simulated the relay cell’s suppressive field are shown separately. Five interneurons recapitulated the structure of the suppressive field. Thus, our work provides proof-of-concept that convergent input from diverse types of interneurons can supply feature-specific inhibition to relay cells in the murine dLGN.

**Figure 7.**
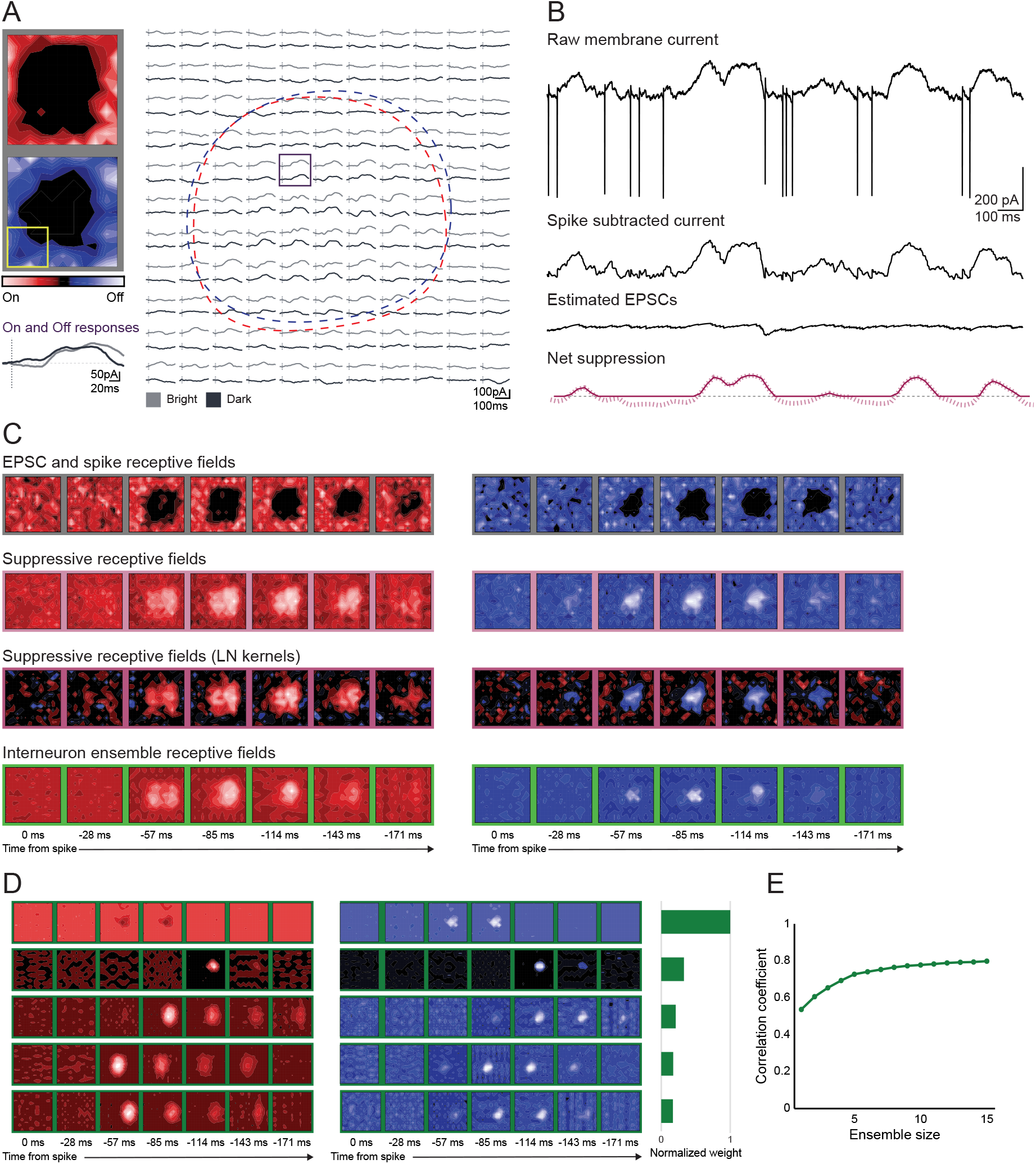
Simulation demonstrating how pooled input from different types of interneurons can generate feature-specific inhibition for an SBC relay cell. Conventions as in Figure 6 except that tSTAs for both On and Off responses are shown for both the relay cell and modeled interneurons.

## DISCUSSION

Local interneurons in thalamus have the power to edit every spike that relay cells transmit from the periphery to cortex. These cells are difficult to sample, however, because they are very small and sparse (Evangelio *et al*., 2018; Jager *et al*., 2021), and cannot be distinguished by spike statistics alone (Pape and McCormick, 1995; Wang *et al*., 2011). Thus, we employed optogenetic approaches in mouse dLGN to distinguish inhibitory from excitatory neurons prior to recording responses to visual stimuli. Our results show that murine interneurons, like neighboring relay cells (Piscopo *et al*., 2013; Suresh *et al*., 2016), have diverse types of receptive fields. The distributions of receptive field structures, but not sizes, differed between the two populations. To evaluate potential contributions of interneurons to the inhibitory component of the relay cell’s receptive field, we turned to computational approaches. Using a library of modeled (Chichilnisky, 2001) interneurons based on our data, and a widely used search algorithm, we showed that a handful of different types of interneurons were able to reproduce the overall spatiotemporal arrangement of suppression in the relay cell’s receptive field. Thus, our work provides proof-of-concept that convergent input from diverse types of interneurons can provide feature-specific inhibition to relay cells.

### Receptive field structure

Before beginning this project, we had hypothesized that murine interneurons might pool retinal input to provide non-selective inhibition to cortex, as is the case for many (Kerlin *et al*., 2010; Liu *et al*., 2009), though not all (Camillo et al., 2018; Runyan et al., 2010), interneurons in murine visual cortex. Rather, we found that the diversity of receptive field shapes for interneurons was comparable to that for relay cells. By contrast, the distribution of receptive field categories differed. For example, there was a greater representation of conventional On-Off cells among interneurons, perhaps a reflection of higher retinal convergence of On and Off pathways. There was also a smaller proportion of center-surround interneurons; we suspect the contributions from ganglion cells with this profile were often masked by convergent input from other retinal afferents (Hammer *et al*., 2015; Maher *et al*., 2023; Morgan *et al*., 2016; Rompani *et al*., 2017; Seabrook *et al*., 2013). Receptive fields characterized by strong suppression (SBC, W3) (Piscopo *et al*., 2013; Suresh *et al*., 2016) were observed only for relay cells in our dataset; recent work using gratings to evaluate direction and orientation selectivity in dLGN shell reports interneurons with SBC-like responses, however (Müllner and Roska, 2023).

Our sample covered much of the anteroposterior and mediolateral extent of the dLGN and included recordings from core and shell. Further, multiconductor probes permitted simultaneous analysis of nearby interneurons and relay cells, which mitigated against regional bias. It is possible that we mistook some interneurons for relay cells. For example, because the LED activated many interneurons at once, these might have inhibited each other and thus prevented ChR2 mediated currents from triggering spikes. Given the small percentage (<10%) of interneurons in dLGN (Evangelio *et al*., 2018; Jager *et al*., 2021), however, this potential confound would have had minimal repercussions.

### Receptive field size

Receptive field sizes of interneurons and relay cells ranged from small and compact to large and amorphous. We had not anticipated finding interneurons with small receptive fields. Previous work in primate (Wilson, 1989; Wilson *et al*., 1996) and cat (Martinez *et al*., 2014) showed that interneurons had receptive fields larger than those of adjacent relay cells. This makes sense because the former vastly outnumber the latter across species (Evangelio *et al*., 2018; Golding et al., 2014; Jager *et al*., 2021; LeVay and Ferster; Montero, 1987) yet must represent the full extent of visual space. Accordingly, more retinal afferents seem to converge onto interneurons than onto relay cells (Maher *et al*., 2023; Morgan *et al*., 2016; Morgan and Lichtman, 2020; Seabrook *et al*., 2013; Van Horn et al., 2000). Further, one might expect that the impact of convergence on receptive field size in mouse would be extreme. The dendritic arbors of murine interneurons extend over far larger regions of retinotopic space (Charalambakis *et al*., 2019; Morgan and Lichtman, 2020; Seabrook *et al*., 2013) than those in carnivore (Hamos *et al*., 1985; Humphrey and Weller, 1988) or primate (Wilson, 1989; Wilson *et al*., 1996). In addition, work *in vitro* shows that active conductances in the dendritic membrane can help to convey strong remote input to the soma (Acuna-Goycolea et al., 2008) and also back propagate action potentials (Casale and McCormick, 2011); this finding is consistent with recent work *in vivo* (Müllner and Roska, 2023) that shows that primary and secondary dendrites share the same direction and orientation preference with the soma when activated with a powerful stimulus.

Why, then, might some interneurons have small receptive fields in the face of high retinal convergence? First, it is possible that only a subset of presynaptic inputs dominates the neural response, as seems to be the case for relay cells (Litvina and Chen, 2017), or that some interneurons only receive input from ganglion cells with small receptive fields, which is possible given the wide range of convergence values for relay cells and interneurons (Jiang et al., 2022; Müllner and Roska, 2023; Rompani *et al*., 2017). Second, some retinal inputs might remain subthreshold. Third, murine interneurons have long thin dendrites (Charalambakis *et al*., 2019) and receive dense retinal input at distal tufts (Morgan and Lichtman, 2020). This remote input might decay passively or because of synaptically induced shunts (Morgan and Lichtman, 2020; Perreault and Raastad, 2006) before reaching the soma. Fourth, thalamic interneurons form dendrodendritic synapses (Morgan and Lichtman, 2020; Sherman, 2004), so subthreshold retinal excitation alone is sufficient to drive postsynaptic inhibition (Cox et al., 1998; Errington et al., 2011; Pressler and Regehr, 2013). Thus, dendritic arbors of interneurons might operate separately from the soma *in vivo,* like retinal amacrine cells (Vaney et al., 2012).

### Stimulus-dependence of feature selectivity

The presence of adaptive receptive fields, those that change categorically as a function of stimulus parameters, suggests another mechanism by which high retinogeniculate convergence is compatible with feature selectivity in dLGN. Examples of adaptive responses included simple shifts from On-Off excitation to sole preference for a single stimulus polarity, to more complex transformations (patterns of suppression also changed but were too subtle to quantify). While stimulus-dependent properties of retinal receptive fields (Tikidji-Hamburyan et al., 2015) likely contribute to this adaptive property, retinogeniculate connectivity and intrinsic inhibition could play a role. That is, different stimuli might engage disparate subsets of convergent retinal inputs on a given target relay cell or interneuron. Although adaptive cells were not common in our sample (15% interneurons, 8% relay cells), they seem virtually absent for center-surround cells in the intact carnivore (Moore et al., 2011; Wang *et al*., 2007) and primate dLGN (Archer et al., 2021) species in which retinogeniculate convergence values are relatively low (Sincich *et al*., 2007; Usrey *et al*., 1999).

### Simulating the suppressive component of the relay cell’s receptive field

With rare exception, relay cells in the main layers of the cat dLGN have center-surround receptive fields with push-pull excitation and inhibition within each subregion. The intuitive explanation for the “pull” is that it derives from local interneurons since these, too, have center-surround receptive fields (Martinez *et al*., 2014; Wang *et al*., 2011; Wilson, 1989). In mouse, however, the situation is complicated. The receptive field structures of murine relay cells (Durand *et al*., 2016; Piscopo *et al*., 2013; Suresh *et al*., 2016) and, as we show here, interneurons, are diverse. Further, the small number of interneurons in mouse (Evangelio *et al*., 2018; Jager *et al*., 2021) and the differences in the distributions of receptive field types between relay cells and interneurons that we report suggested that various types of interneurons converge onto a single postsynaptic target.

To explore whether or not convergent input from distinct interneurons could, in principle, simulate shapes of suppressive receptive fields recorded from relay cells, we developed a simple computational framework. First, we constructed models of receptive fields obtained from interneurons, and proceeded with the assumptions that all types of interneurons are available to all relay cells, and that different interneurons within the same class vary slightly in latency and tile the visual field. We began each simulation by cycling through the pool of modeled interneurons to choose the one that contributed most to the suppressive component of a given relay cell’s response, and then iterated this process to select additional inputs. In this fashion, we were able to identify groups of interneurons whose aggregated input reproduced the overall shape of the relay cell’s suppressive receptive field.

Altogether, our approach shows that local interneurons in mouse dLGN have diverse types of receptive fields and demonstrates how convergent input from these neurons can, in principle, explain the patterns of inhibition recorded from relay cells. These results provide a scheme for how inhibitory inputs might be combined in murine thalamic nuclei (e.g., somatosensory or auditory) with even smaller (<6%) proportions of interneurons (Jager *et al*., 2021). Our results also relate to species with far higher (∼25-35%) proportions of thalamic interneurons (LeVay and Ferster; Montero and Zempel, 1986). For example, there is compelling evidence for convergence of On and Off inhibition in cat and monkey dLGN via a scheme in which relay cells receive excitatory as well as inhibitory input initiated by the same retinal afferent (Guillery, 1969; Hamos *et al*., 1985; Wilson, 1989) in addition to connections from interneurons of the opposite sign (Wang *et al*., 2011). Perhaps future experiments will specify patterns of inhibition that relay cells receive by imaging presynaptic boutons *in vivo*, as has been accomplished for retinal terminals that supply dendrites in mouse dLGN (Liang et al., 2018). Last, we note that our study was limited to exploring contributions of local interneurons. In the future, it will be important to include input from the thalamic reticular nucleus, an inhibitory structure that receives input from relay cells and synapses on them in return (Born et al., 2021; Campbell et al., 2020; Soto-Sánchez et al., 2017; Vaingankar *et al*., 2012).

## Author contributions

ASG and JAH designed the research plan; ASG, SA, and YM conducted experiments involving optogenetically labeled cells; VS and UMC contributed whole-cell recordings from relay cells; AG developed quantitative analyses; YM and FTS built model frameworks to link interneuron output to relay cell suppression; YS, UMC and FTS contributed analysis pipelines. ASG, YM, and JAH wrote the manuscript and all other authors provided comments and approved the final text.

## Acknowledgments

The authors thank Martha Bickford. for reading the manuscript and NIH R01 EY09395 to JAH and NSF IIS 1718991 and NIH 1R01 EB026955 to FTS

The authors declare no competing interests.

---

**Figure S1.**
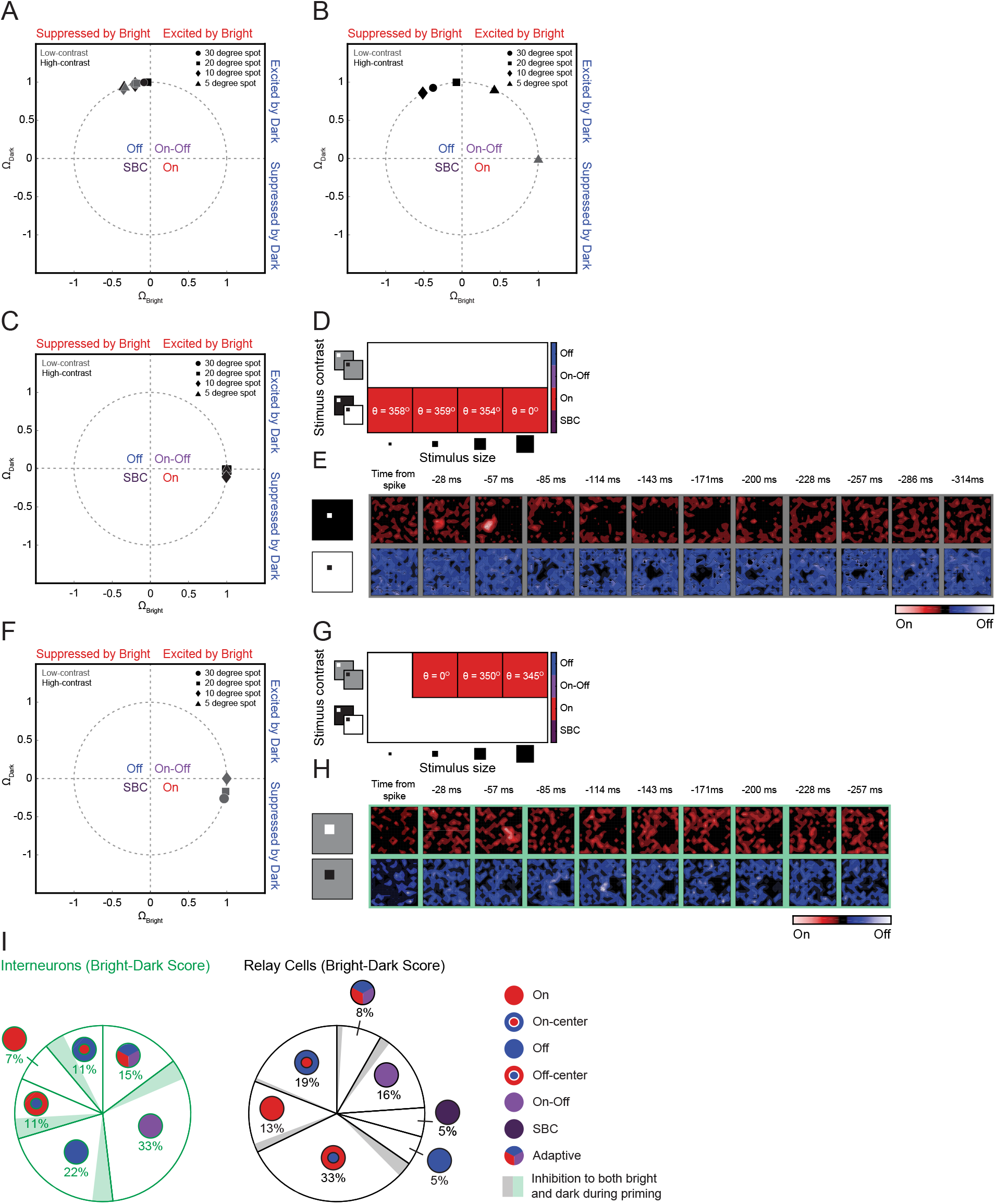
Bright-Dark Polarity Scores for cells with prolonged suppression to both bright and dark stimuli during priming phase. (A) Bright-Dark polarity scores for the cell represented in Figures 4C and 4D shown on a unit circle. Triangles, diamonds, squares, and circles mark data points calculated from responses to 5°, 10°, 20°, and 30° squares, respectively; black and gray markers are data points calculated from responses to full and half-contrast stimuli, respectively. Inset shows scores at magnified view. (B) Bright-Dark polarity scores for Figures 4E and 4F, conventions as in A. (C-E) Example of an On-relay cell whose response included unusually prolonged suppression to both bright and dark stimuli during the priming phase. (C) Bright-Dark polarity scores for a relay cell shown along a unit circle (conventions as in A), and (D) as the angle θ. (E) The tSTAs for On (top) and Off (bottom) stimuli computed from responses to 5° squares, full contrast sparse noise; prolonged suppression during the priming phase is especially prominent for the Off response. (F-H) An example of an interneuron with prolonged suppression during the priming phase. Conventions as for C-E except the stimulus used for the tSTAs was different; 20° squares, half-contrast sparse noise. (I) Pie-charts of different types of receptive fields for interneurons and relay cells modified from Figure 4I to include cells with (qualitatively assessed) prolonged suppression during the priming phase.

**Figure S2.**
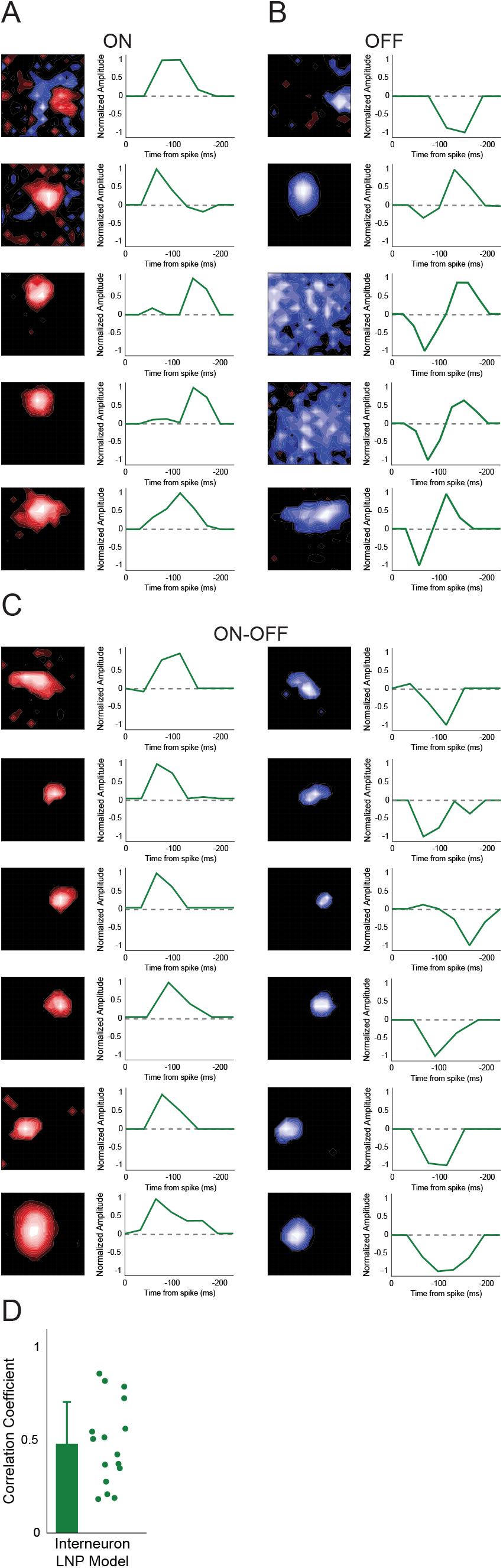
LNP models for 16 interneurons. Spatial (left) and temporal (right) filters for On interneurons (A), Off interneuron and (B), and On-Off interneurons. (D) Correlation coeffcients for LNP models and tSTAs.

## Notes

### Competing Interest Statement

The authors have declared no competing interest.

